# Translocation of gut commensal bacteria to the brain

**DOI:** 10.1101/2023.08.30.555630

**Authors:** Manoj Thapa, Anuradha Kumari, Chui-Yoke Chin, Jacob E. Choby, Fengzhi Jin, Bikash Bogati, Daniel M. Chopyk, Nitya Koduri, Andrew Pahnke, Elizabeth J. Elrod, Eileen M. Burd, David S. Weiss, Arash Grakoui

## Abstract

The gut-brain axis, a bidirectional signaling network between the intestine and the central nervous system, is crucial to the regulation of host physiology and inflammation. Recent advances suggest a strong correlation between gut dysbiosis and neurological diseases, however, relatively little is known about how gut bacteria impact the brain. Here, we reveal that gut commensal bacteria can translocate directly to the brain when mice are fed an altered diet that causes dysbiosis and intestinal permeability, and that this also occurs without diet alteration in distinct murine models of neurological disease. The bacteria were not found in other systemic sites or the blood, but were detected in the vagus nerve. Unilateral cervical vagotomy significantly reduced the number of bacteria in the brain, implicating the vagus nerve as a conduit for translocation. The presence of bacteria in the brain correlated with microglial activation, a marker of neuroinflammation, and with neural protein aggregation, a hallmark of several neurodegenerative diseases. In at least one model, the presence of bacteria in the brain was reversible as a switch from high-fat to standard diet resulted in amelioration of intestinal permeability, led to a gradual loss of detectable bacteria in the brain, and reduced the number of neural protein aggregates. Further, in murine models of Alzheimer’s disease, Parkinson’s disease, and autism spectrum disorder, we observed gut dysbiosis, gut leakiness, bacterial translocation to the brain, and microglial activation. These data reveal a commensal bacterial translocation axis to the brain in models of diverse neurological diseases.

---

Recent evidence suggests a strong correlation between gut dysbiosis and neurodegenerative diseases such as Alzheimer’s disease (AD), Parkinson’s disease (PD), and autism spectrum disorder (ASD) <u>^1-4^</u>. Gut bacteria have been shown to affect neurodegeneration indirectly via secreted metabolites and toxins; however, it is unclear if they have a direct impact on the brain <u>^1,5-8^</u>. Here, utilizing several murine models of gastrointestinal and neurodegenerative diseases, we observed gut commensal bacterial translocation directly to the brain, associated with markers of neuroinflammation and neural protein aggregate formation.

## Gut bacteria translocate to the brain

We previously observed alteration of gut microbiome composition in a mouse model of cholestatic liver disease (multidrug resistance gene 2 knockout, *Mdr2^-/-^*) <u>^9^</u>. Here, we tested how diet alteration might impact the gut microbiome in this model, by feeding mice an atherogenic, high-fat Paigen diet. Total bacterial colony forming units (CFU) were comparable in fecal samples and the ileum in *Mdr2^-/-^* mice fed control or Paigen diet. However, after 8 days of Paigen diet, *Mdr2^-/-^* mice exhibited dysbiosis characterized by marked enrichment of *Staphylococcus*, *Bacteroides* and *Akkermansia*, and a reduction of gut commensal *Lactobacilli* in the feces, as compared with mice fed a control diet (**Fig. 1a**). This dysbiosis correlated with increased gut leakiness in mice fed Paigen diet (**Fig. 1b**). To determine whether the intestinal leakiness in these Paigen diet-fed mice might lead to bacterial dissemination out of the intestine, we cultured the blood and plated organs such as the lung, heart, kidney, spleen, and brain. While we did not observe bacteria in the blood (sensitivity <5 CFU) or other systemic organs (**Supplementary Fig. S1**), we were surprised to observe bacteria in the brains of Paigen diet-fed, but not control diet-fed mice (**Fig. 1c-g**). Matrix-assisted laser desorption/ionization time-of-flight (MALDI-TOF) mass spectrometry identified the bacteria isolated from the brains as *Staphylococcus xylosus* (*S. xylosus*), a mammalian commensal bacterium, consistent with the enrichment of *Staphylococci* in the feces and ileum of Paigen-fed *Mdr2^-/-^* mice (**Fig. 1h-k**).

**Figure 1.**
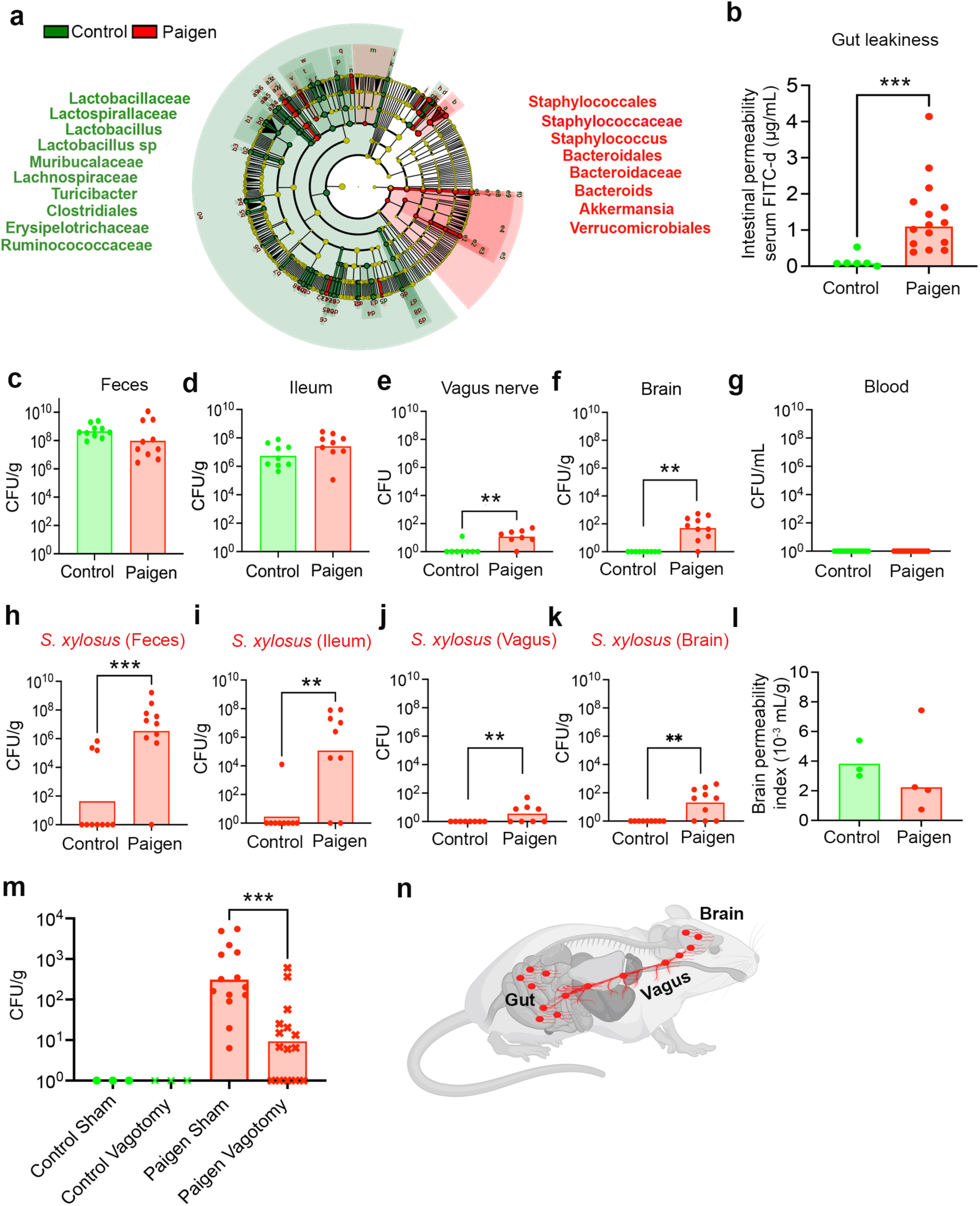
Gut bacteria translocate to the brain via vagus nerve. Paigen diet was fed to 12-week-old *Mdr2^-/-^*mice for 8 days. **a)** Fecal pellets collected at day 8 were processed for 16S genomic sequencing. Cladogram representation showing the phylogenetic relationship between bacterial families between control diet (green) and Paigen diet (red) groups are shown. **b**) Intestinal permeability was determined by using fluorescein isothiocyanate-dextran (4 Kda) oral gavage assay. **c-g**) Total colony forming units (CFU) of bacteria (**c**) per gram of feces, (**d**) per gram of ileum, (**e**) vagus nerve, (**f**) per gram of brain and (**g**) per ml of blood specimens are shown. **h-k**) CFU of *Staphylococcus xylosus* in (**h**) feces, (**i**) ileum, (**j**) vagus nerve and (**k**) brain are shown. **l**) Brain permeability index of control diet and Paigen diet fed mice are shown. **m**) Total CFU/g of brain following unilateral right cervical vagotomy of mice are shown. **n**) A schematic representation of a commensal bacterial translocation axis to vagus nerve and the brain is demonstrated in paigen diet-fed mice (generated from Biorender). Data presented from minimum 2 independent experiments (n=3– 5/group). Statistical significance was determined by Mann Whitney test or one-way analysis of variance (ANOVA) where applicable. \*\**P*<0.01, \*\*\**P*<0.001. Bars reflect the geometric mean.

## Gut bacteria translocate to the brain via the vagus nerve

It was unclear how *S. xylosus* translocated to the brain and why this localization was specific and not also observed in systemic organs. The localization of bacteria in the brains of Paigen diet-fed mice was not due to increased blood-brain barrier permeability, which was comparable to mice fed the control diet (**Fig. 1l**). As the vagus nerve connects the gut and the brain, we tested the vagus nerve itself for the presence of bacteria. We isolated *S. xylosus* from the vagus nerve (**Fig. 1j**) but not the spinal cord (**Supplementary Fig. S1e**), indicating that the localization of these bacteria to the vagus nerve was specific. To test whether the vagus nerve could serve as a route of bacterial translocation to the brain, we next disrupted the right cervical vagus nerve by vagotomy <u>^10,11^</u>. Vagotomized and sham operated mice were subsequently fed Paigen diet. We observed that vagotomized mice fed Paigen diet harbored ∼20-fold lower levels of bacteria in the brain as sham controls (some vagotomized mice had no detectable bacteria in the brain), whereas brains from both vagotomized and sham controls fed conventional diet did not contain bacteria (**Fig. 1m**). Since vagotomy was done only on the right cervical nerve (bilateral cervical vagotomy would be fatal), the left cervical vagus nerve remained intact and therefore could have been used as an alternative route by which bacteria translocated to the brain, since we observed that bacteria can localize to both the right and left branches (**Supplementary Fig. S2**). Taken together, these data reveal a pathway by which commensal gut bacteria can translocate to the brain via the vagus nerve (**Fig. 1n**). While we cannot rule out the contribution of other routes, these data indicate that commensal translocation can occur via the vagus nerve and in the absence of any observable blood-brain barrier permeability.

## Intestinal microbiome composition dictates which bacteria translocate to the brain

To further explore the link between intestinal bacteria and those that translocate to the brain, we perturbed the microbiome by treating mice with a cocktail of antibiotics (vancomycin, 500mg/L; ampicillin 1g/L; neomycin, 1g/L; and metronidazole, 1g/L) in their drinking water during Paigen diet feeding. While *Staphylococcus sp.* were highly enriched in the ileum and feces of Paigen diet-fed mice, the addition of antibiotic treatment (Abx) reduced the levels of *Staphylococci* and led to the enrichment of *Paenibacillus cineris* (*P. cineris*) (**Supplementary Fig. S3**). Of note, antibiotic treatment did not further exacerbate the intestinal permeability caused by Paigen diet (**Supplementary Fig. S3a**). We again observed a correlation between the bacteria in the gut and those found in the vagus nerve and brain, as Paigen diet-fed, antibiotic-treated mice harbored *P. cineris* or *Staphylococci* in their vagus nerve and brain (**Supplementary Fig. S3b-i**). Similar to the previous data with *S. xylosus* in Paigen diet-fed mice without antibiotics, we could not detect *P. cineris* or *Staphylococci* in the blood, systemic organs, or the spinal cord. Further, Paigen diet-fed and antibiotic-treated mice exhibited no increase in blood-brain barrier permeability. These data further implicate the gut microbiome as the source of brain-localizing bacteria, demonstrating that a perturbation that changes the composition of the microbiome (i.e., diet and antibiotic use) analogously changes the bacteria that localize to the brain.

To determine whether commensal bacterial translocation to the brain occurs in mouse backgrounds other than *Mdr2^-/-^*, we fed wild-type C57BL/6 mice Paigen diet for 2 weeks. Paigen diet altered the microbiome composition and caused an increase in gut leakiness in these mice (**Fig. 2a-d, Supplementary Fig. S4**). C57BL/6 mice exhibited enriched *S. xylosus* and *Enterococcus faecalis* (*E. faecalis*) in their feces and ilea (**Supplementary Fig. S4)**. Correspondingly, we detected both *S. xylosus* and *E. faecalis* in the brains of Paigen diet-fed C57BL/6 mice, further demonstrating the specific link between gut bacterial composition and the bacteria in the brain (**Supplementary Fig. S4**). Next, to determine whether we could exogenously alter the bacteria that reach the brain, we first disrupted gut bacterial composition by treating C57BL/6 mice with a cocktail of antibiotics. We subsequently gavaged *Enterobacter cloacae* (*E. cloacae*) into the antibiotic-treated mice and fed them Paigen diet (**Fig. 2e**). *E. cloacae* were detected in the gut on day 5 after gavage (**Fig. 2f, g**). On day 8 after gavage, *E. cloacae* were detected in the brain of the C57BL/6 mice (**Fig. 2h**), further confirming that the bacteria found in the brain are specific to those colonizing the gut. Taken together, these data show that bacteria can translocate to the brain of multiple genetic types of mice including wild-type mice, that numerous types of bacteria can translocate to the brain even simultaneously, and that in all cases studied thus far, the bacteria detected in the brain are also found in the intestine.

**Figure 2.**
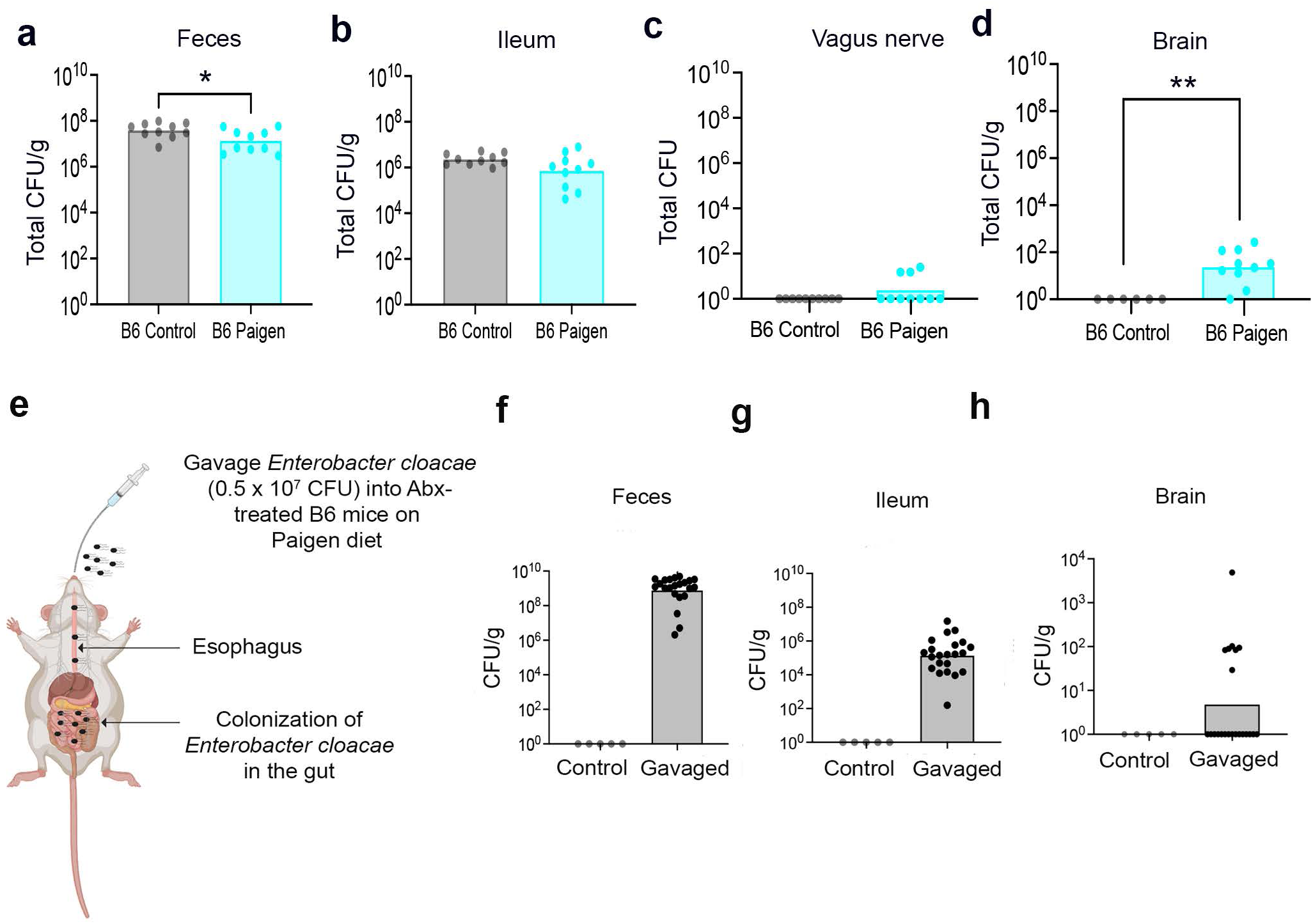
Gut microbiome composition dictates which bacteria translocate to the brain. CFU of bacteria in (**a**) feces, (**b**) ileum, (**c**) vagus nerve and (**d**) brain of C57BL/6 (B6) mice fed Paigen diet are shown. **e**) *Enterobacter cloacae* (5×10^7^ CFU) were gavaged into Abx-treated B6 mice fed Paigen diet (generated from Biorender). **f-h**) CFU of *Enterobacter cloacae* in (**f**) feces, (**g**) ileum and (**h**) brain tissues were determined. Data presented from minimum 2 independent experiments (n=3–5/group). Statistical significance was determined by Mann Whitney test or one-way analysis of variance (ANOVA) where applicable. \**P*<0.05. Bars reflect the geometric mean.

## Bacterial localization to the brain triggers microglial activation and neural protein aggregates

Neuroinflammation is now recognized as a major component of many neurodegenerative and neurodevelopmental diseases. Since bacteria are known to be inflammatory, we next set out to determine if brains harboring translocated commensal gut bacteria exhibit hallmarks of neuroinflammation. Microglia are phagocytic cells in the brain that promote inflammation. These cells exhibit a highly branched resting morphology in healthy states but a rounded, smaller amoeboid morphology in disease states. Ionized calcium-binding adapter molecule (IBA-1) is a microglia marker <u>^12-14^</u>, whose staining can be used to calculate a measure of microglia morphology and activation status called the ramification index (**Fig. 3a**). The microglia ramification index was two-fold increased in the brains of Paigen diet-fed *Mdr2^-/-^* mice (**Fig. 3b**), which we showed harbor translocated commensal bacteria in the brain (**Fig. 1f, k**), indicating a smaller, rounded amoeboid morphology characteristic of activated cells that are associated with neuroinflammation.

**Figure 3.**
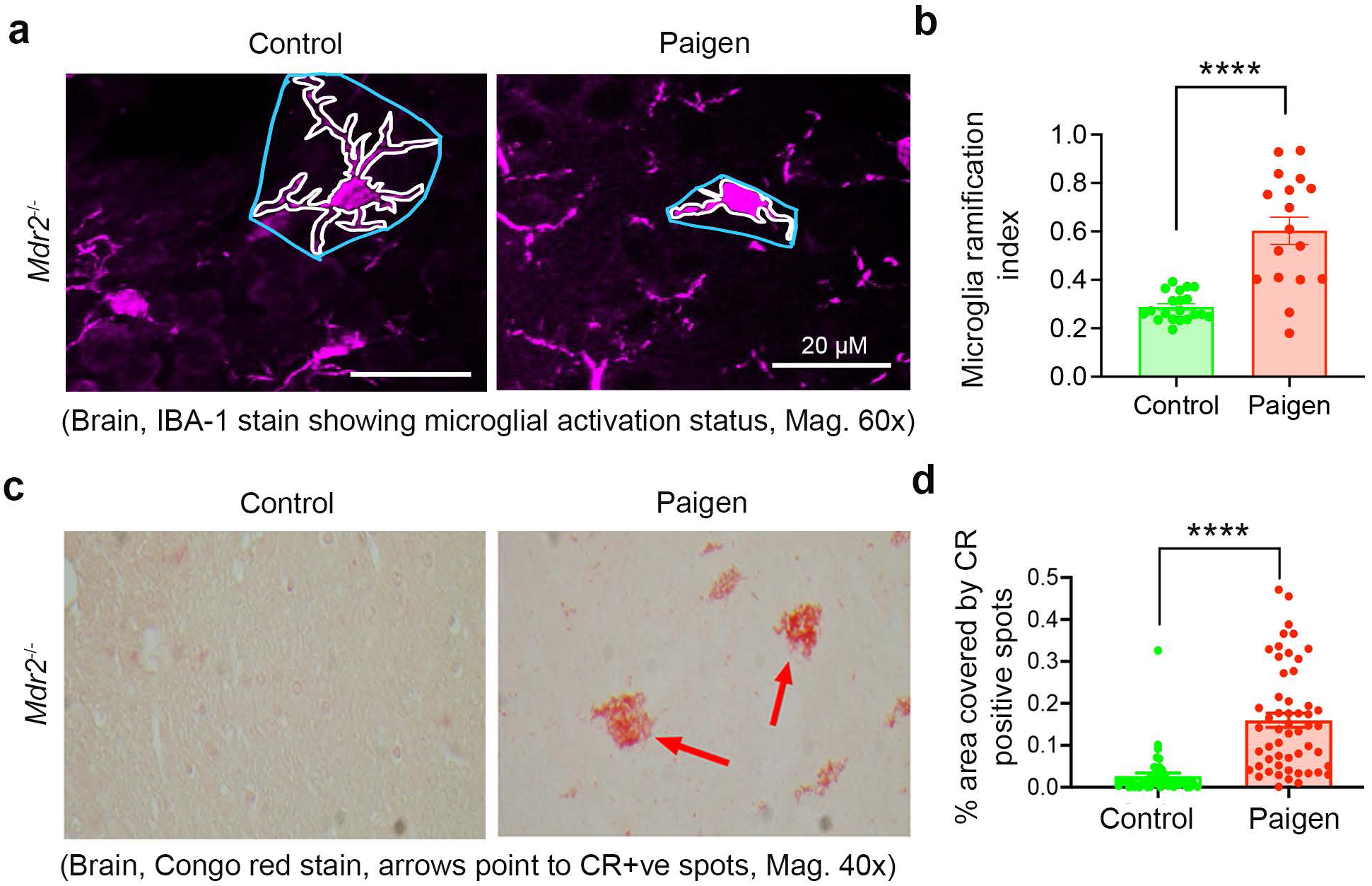
Bacterial localization to the brain triggers neuroinflammation and protein aggregation. **a**) Representative immunofluorescence IBA-1 staining of brain sections showing microglial activation status in control diet and Paigen diet-fed *Mdr2^-/-^*mice. **b**) Microglial ramification index in control diet and Paigen diet-fed *Mdr2^-/-^* mice are shown. **c**) Representative Congo red stained brain sections from control diet and Paigen diet-fed *Mdr2^-/-^* mice. **d**) Percent area covered by Congo red positive spots per field of brain sections from control diet and Paigen diet-fed *Mdr2^-/-^* mice are shown. Data presented from minimum 2 independent experiments (n=3–5/group). Statistical significance was determined by Mann Whitney test or one-way analysis of variance (ANOVA) where applicable. \*\*\*\**P*<0.0001. Bars reflect the geometric mean.

The aggregation of proteins in the brain is considered one of the hallmark features of neurodegenerative disorders <u>^15,16^</u>. To investigate whether the localization of gut bacteria to the brain is associated with neural protein aggregates, we performed Congo red staining of brain sections from Paigen diet-fed *Mdr2^-/-^* mice. We observed localized Congo red-positive staining in the brains of Paigen diet-fed *Mdr2^-/-^* mice, indicating the presence of protein aggregates (**Fig. 3c-d**). Taken together, these data identify a correlation between the localization of gut bacteria in the brain, microglial activation, and neural protein aggregation.

## Restoration of conventional diet reverses bacterial localization, microglial activation, and neural aggregate formation in the brain

Because *Mdr2^-/-^* mice fed a Paigen diet developed intestinal permeability, translocation of commensal gut bacteria to the brain, microglial activation, and neural protein aggregation, we tested whether these phenotypes could be reversed upon restoration of a conventional diet. To do so, we fed Paigen diet to *Mdr2^-/-^* mice for 7 days and confirmed they exhibited the aforementioned phenotypes. We subsequently reversed their diet back to normal rodent chow and tested their phenotypes at days 14 (D14) and 28 (D28). Restoration of a conventional diet decreased intestinal leakiness by four-fold on D14 and D28 (**Fig. 4a**). Consistent with these observations, with only two exceptions, CFU of *S. xylosus* were below the detectable limit in both the ileum and the brain on D14 (**Fig. 4b, c**). The reduction in *S. xylosus* in the brain correlated with a reduced microglia ramification index following diet reversal on D14, indicating that microglial activation was reversed (**Fig. 4d, e**). At D14, protein aggregates were not reduced in number or area in the brain, but by D28 the percent area covered by aggregates dropped by almost 5-fold (**Fig. 4f, g**). Thus, the reduction in neural protein aggregates was preceded by the reduction in *S. xylosus* in the brain and reduced microglia activation. These data, in at least one model, highlight the reversibility of bacterial localization to the brain and neural protein aggregates, and suggest that the reduction of dysbiosis and intestinal permeability may be important attributes of effective therapies to treat neurodegenerative and neurodevelopmental diseases.

**Figure 4.**
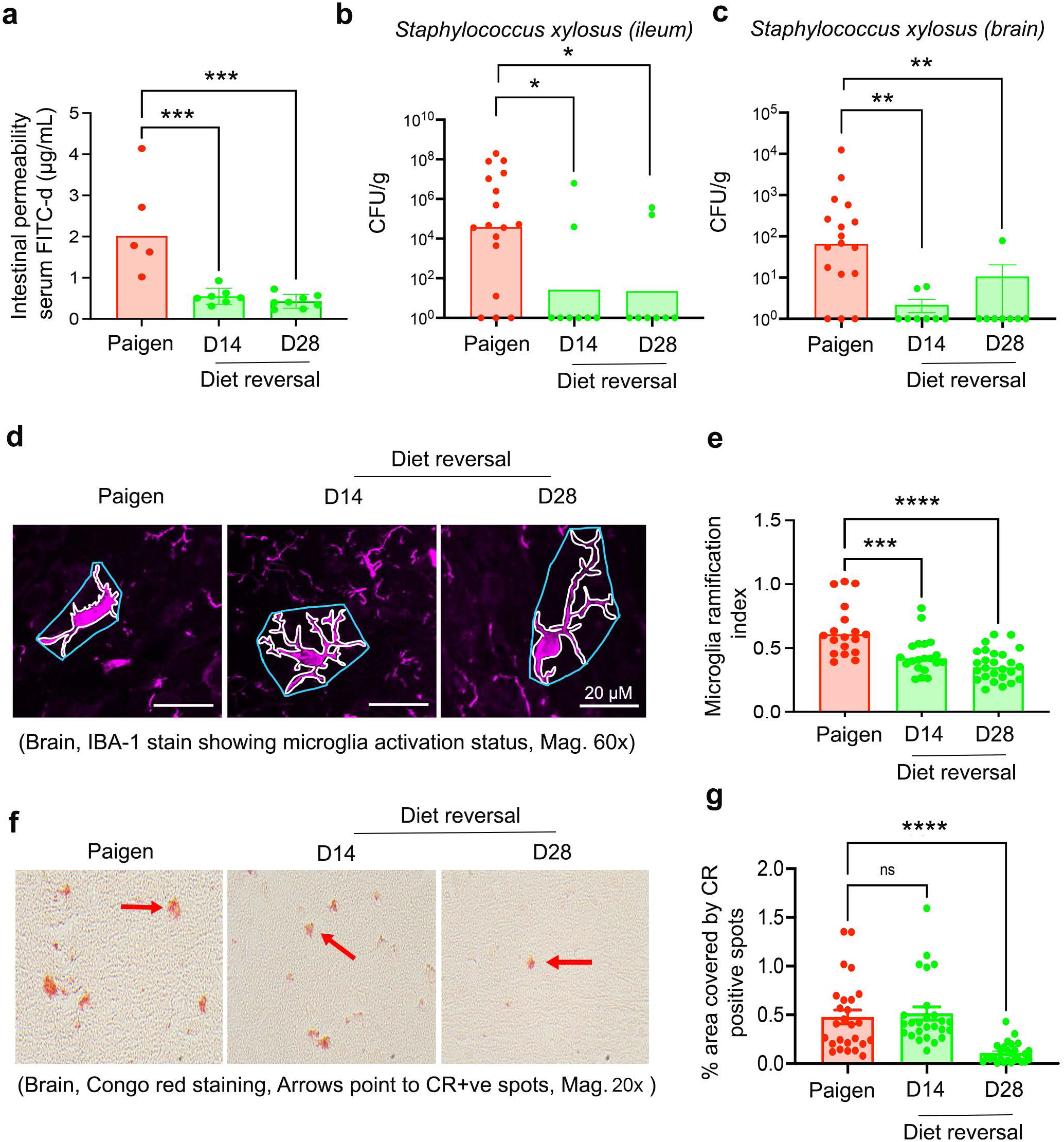
Diet reversal reduces gut permeability, bacterial localization to the brains, plaque formation, and neuroinflammation. Paigen diet was fed to 12-week-old *Mdr2^-/-^*mice for 8 days (“Paigen” group), before reversal to conventional diet for 14 (“D14”) or 28 (“D28”) days. **a)** Intestinal permeability was determined by using fluorescein isothiocyanate-dextran (4 Kda) oral gavage assay. **b-c**) Colony forming units (CFU) of *Staphylococcus xylosus* (**b**) per gram of ileum and (**c**) per gram of brain are shown. **d**) Representative immunofluorescence IBA-1 staining of brain sections showing microglial activation status in Paigen diet-fed *Mdr2^-/-^* mice or those switched to conventional diet. **e**) Microglial ramification index in Paigen diet-fed *Mdr2^-/-^* mice or mice reversed to conventional diet is shown. **f**) Representative Congo red stained brain sections from Paigen diet-fed *Mdr2^-/-^* mice or mice subsequently switched to conventional diet for 14 or 28 days. **g**) Percent area covered by Congo red positive spots per field of brain sections from control diet and Paigen diet-fed *Mdr2^-/-^* mice are shown. Data presented from minimum 2 independent experiments (n=3–5/group). Statistical significance was determined by Mann Whitney test or one-way analysis of variance (ANOVA) where applicable. \**P*<0.05, \*\**P*<0.01, \*\*\**P*<0.001, \*\*\*\**P*<0.0001. Bars reflect the geometric mean.

## Commensal bacterial localization in the brains of mouse models of neurodegenerative diseases

Recent evidence implicates the disruption of the gut microbiome and gut permeability as being associated with neurodegenerative diseases such as Alzheimer’s disease (AD), Parkinson’s disease (PD) and neurodevelopmental diseases such as autism spectrum disorder (ASD) <u>^2,17-19^</u>. We investigated mouse models for each disease (AD, APP/PS1 WSB.Cg-Tg(APPswe,PSEN1dE9)85Dbo/How; PD, LRRK2 knock-in (B6.Cg-Lrrk2^tm1.1Hlme^/J; and ASD, BTBR T+ Itpr3^tf^/J BTBR) to determine if dysbiosis and intestinal permeability might be linked to brain translocation of bacteria in these mice. First, we quantified the microbiota composition in these mice. APP/PS1 mice exhibited enriched *Streptococcus*, *Clostridium*, *Roseburia* and *Tyzzerella* in comparison to wild-type mice, while LRRK2 mice exhibited marked enrichment of *Staphylococcus*, *Eubacterium, Enterorhabdus, Coriobacteria* and *Flavonifractor* (**Fig. 5a**). BTBR mice showed a broader enrichment of bacteria compared to wild-type mice, including *Firmicutes, Corynebacterium, Anaeroplasma, Monoglobus, Fusimonas, Murimonas, Tyzzerella, Lachnospiraceae, Erysipelotrichaceae, Enterococcus, Staphylococcus* and *Robinsoniella*. In line with previous observations, we observed the disruption of the intestinal barrier (i.e., gut leakiness) to be a common phenomenon in these mouse models of AD, PD, and ASD (**Fig. 5b)**.

**Figure 5.**
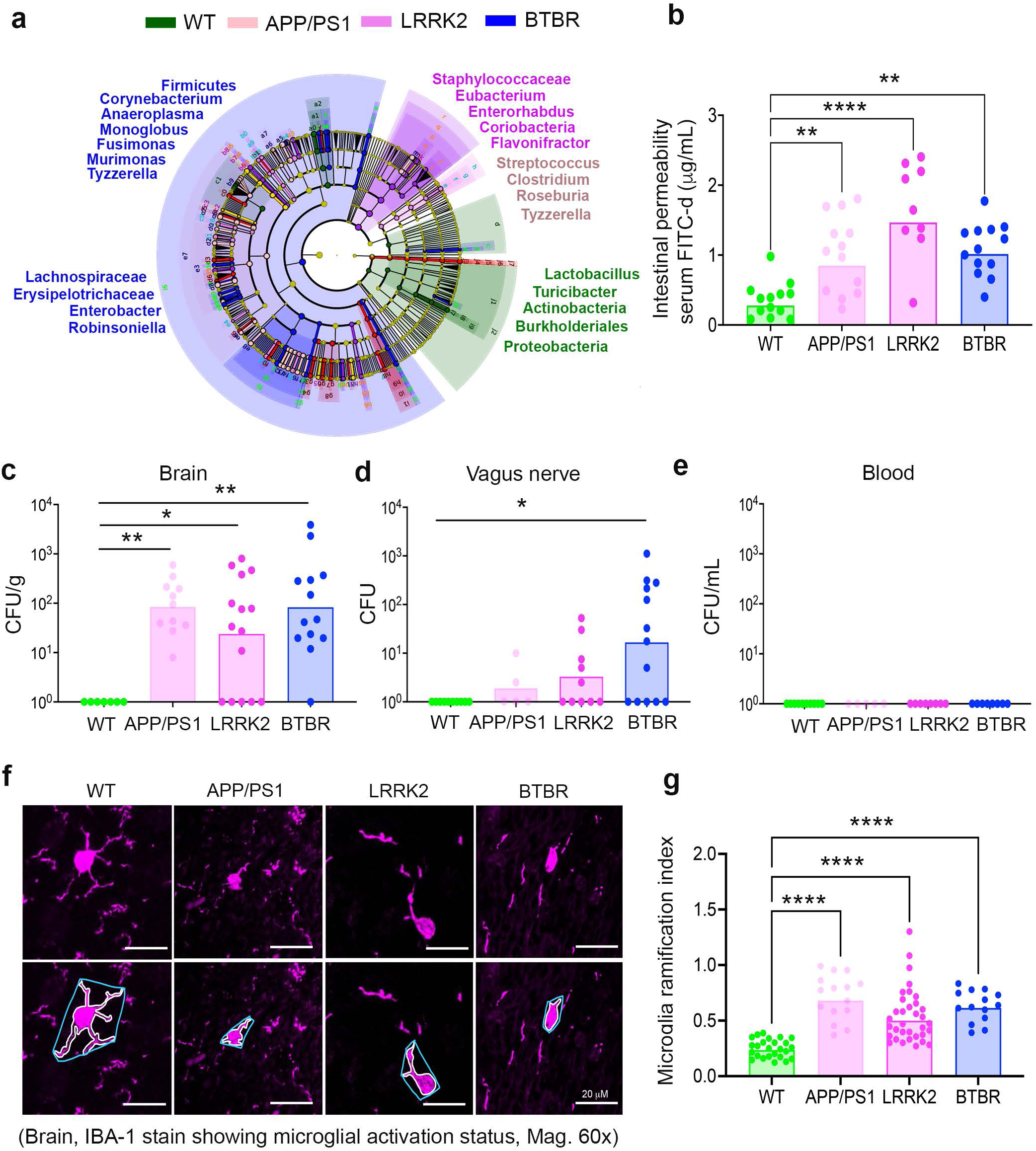
Commensal bacterial localization in the brains of mouse models of neurodegenerative diseases. **a)** Fecal pellets collected from WT, APP/PS1, LRRK2 and BTBR mice from 8-15 wks old mice were processed for 16S genomic sequencing. Cladogram representation showing the phylogenetic relationship between bacterial families between WT, APP/PS1, LRRK2 and BTBR mice groups are shown. **b**) Intestinal permeability was determined by using fluorescein isothiocyanate-dextran (4 Kda) oral gavage assay. **c-e**) Total colony forming units (CFU) of bacteria in (**c**) brain, (**d**) vagus nerve, and (**e**) blood specimens are shown. **f**) Representative immunofluorescence IBA-1 staining of brain sections showing microglial activation status in WT, APP/PS1, LRRK2 and BTBR mice. **g**) Microglial ramification index in WT, APP/PS1, LRRK2 and BTBR mice are shown. Data presented from minimum 2 independent experiments (n=3– 5/group). Statistical significance was determined by Mann Whitney test or one-way analysis of variance (ANOVA) where applicable. \**P*<0.05, \*\**P*<0.01, \*\*\*\**P*<0.0001. Bars reflect the geometric mean.

Gut commensal bacteria were detected in the brain as well as the vagus nerve in these murine models of AD, PD, and ASD (**Fig. 5c, d**). Further, the bacteria detected in the brain and the vagus nerve in these strains were 100% matched with those that were detected in the feces and ileum. Bacterial culture of tissues and blood specimens (**Fig. 5e**) found no detectable bacterial growth in *in vitro* culture conditions among these strains, and brain permeability indexes were not elevated compared to controls. In addition, each of these mice exhibited increased microglial activation as measured by the microglia ramification index (**Fig. 5f, g**). Taken together, these findings provide evidence that gut dysbiosis and altered gut permeability leading to gut commensal bacterial translocation to the brain occur in models of both neurodegenerative and neurodevelopmental diseases.

## Discussion

Using diverse murine models of gastrointestinal disease, neurologic disease, and diet modulation, we demonstrate that commensal gut bacteria can translocate to the brain. The direct link between intestinal bacteria and those isolated in the brain is supported by numerous data; 1) each bacterium isolated from the brain is known to colonize the intestine, 2) in all models tested, the bacteria isolated from the brain were also detected in fecal or ileal samples, 3) modulation of the composition of the intestinal microbiome with antibiotics correspondingly changed the type of bacteria that localized to the brain, 4) intestinal colonization of mice with exogenously delivered bacteria not otherwise detected in the brain led to isolation of those exogenous bacteria from the brain, and 5) in all models in which we observed commensal bacteria in the brain, we also observed an increase in intestinal permeability, providing a rationale for how the bacteria could have escaped the intestine.

The translocation of commensal bacteria to the brain occurred in the absence of permeability of the blood-brain barrier, detectable bacteria in the blood, or broader dissemination of the bacteria to other systemic organs. Rather, our data indicate that the vagus nerve can serve as an anatomical conduit that facilitates, at least in part if not wholly, this translocation; 1) bacteria were isolated from both sides of the cervical vagus nerve, 2) the bacteria in the vagus nerve matched those isolated from the brain and were also detected in the intestinal microbiome, and 3) unilateral cervical vagotomy reduced the number of bacteria in the brain. Of note, we suggest that the residual bacteria isolated from the brain after unilateral cervical vagotomy likely translocated through the intact side of the vagus nerve, although we cannot rule out at least some translocation via a different pathway.

It is important to emphasize that we have observed relatively low-level bacterial localization to the brain (between 1 and ∼300 bacterial cells) that is distinct from an acute, fulminant brain infection. Nevertheless, this low bacterial load is associated with microglial changes associated with neuroinflammation, as well as the formation of neural protein aggregates (**Fig. 3a-d**, **Fig. 5f, g**). Further, we observed these phenotypes in young mice (∼8-15 weeks old), long before the characteristic disease-related changes that occur in some of the mouse models employed here. Therefore, the data suggest that commensal bacterial translocation to the brain is an early event and could even be an initiating trigger for microglial changes associated with neuroinflammation and neural protein aggregate formation, leading to certain neurodegenerative and neurodevelopmental diseases. Since we observe bacterial localization to the brain in multiple models of neurodegenerative and neurodevelopmental disease, this raises the surprising potential for a common link between certain diverse neurological diseases.

These data show how genetic and environmental factors, alone or in combination, can contribute to physiological alterations leading to translocation of gut bacteria to the brain and subsequent hallmarks of neurological disease. It is intriguing that in at least one model, reversal of the environmental condition promoting bacterial translocation (i.e., high-fat diet), led to a reversal of intestinal dysbiosis, permeability, bacterial localization to the brain, microglial activation, and a return to baseline levels of neural protein aggregates in the brain. This suggests that at least some of the processes leading to neuroinflammation and protein aggregate formation are dynamic and raise the potential for future therapeutic interventions capable of reversing these phenomena.

## Materials and methods

### Mouse strains

C57BL/6J (000664), FVB/NJ (001800) mice (wild-type controls) and *Mdr2^-/+^*(FVB.129P2-Abcb4*^tm1Bor^*/J) mice were obtained from the Jackson Laboratory (ME, USA) and housed in a specific pathogen-free environment, per Emory University, NIH, and IACUC guidelines. *Mdr2^-/-^* mice, *Mdr2^-/+^*mice, and control mice were generated by backcrossing littermates until desired true breeding lines were established and confirmed by PCR. BTBR T+ Itpr3^tf^/J (JAX 002282), APP/PS1 WSB.Cg-Tg(APPswe,PSEN1dE9)85Dbo/How (JAX 025970), and LRRK2 knockin (B6.Cg-Lrrk2^tm1.1Hlme^/J (JAX 030961) mice were purchased from the Jackson Laboratory and housed in a specific pathogen free environment per Emory University, NIH and IACUC guidelines. A standard chow (control; Labdiet 5001) or Paigen’s Atherogenic Rodent diet (Research Diets D12336), was fed to *Mdr2^-/-^* male and female mice for 8–12 days. In some experiments, mice were given a cocktail of antibiotics (neomycin sulfate, 1g/L; ampicillin 1g/L; vancomycin 500g/L; and metronidazole 1g/L) in drinking water, while on HFD treatment.

### Intestinal permeability assay

Mice were fasted for 8-10 hours and gavaged with 100 μL of 160 mg/mL FITC-dextran (4 kDa) dissolved in PBS. Blood was collected from mice 4 hours post-gavage by submandibular bleed, and serum was collected after centrifugation. Dilutions of the remaining FITC dextran solution ranging from 15.6 ng/mL to 8000 ng/mL were made for the standard curve. Serum was diluted from 1:2 to 1:10 depending on the volume of serum recovered. The fluorescence of diluted serum and dilutions for the standard curve was measured in a fluorimeter after excitation of the samples at 488 nm and emission at 521 nm.

### Blood-brain barrier permeability index

The blood-brain barrier permeability index (BPI) was calculated using the FITC tracer as described previously <u>^20^</u>. Briefly, the mice were injected intraperitoneally with 100 μl of 2mM FITC dextran (3KDa) solution in PBS. After 20 minutes, mice were euthanized, and blood was collected by cardiac puncture, followed by intracardiac perfusion with 20-40 mL of PBS. The brain was collected in 1 mL PBS and homogenized. The homogenate was centrifuged at 15,000g for 20 minutes, and the fluorescence intensity (F.I) of FITC in the supernatant, along with F.I. of FITC in serum, was measured using a fluorescence plate reader. The F.I. per gram of the brain was determined by dividing the F.I. of the brain homogenate by the brain weight (normalized brain F.I.). Similarly, F.I. per mL of serum was calculated by dividing the F.I. of the serum by the serum volume used for fluorescence calculation (normalized serum F.I.). The BPI was calculated by dividing the normalized brain F.I. by normalized serum F.I.

### Blood culture

Mice were euthanized by CO_2_ asphyxiation. Blood was collected terminally from the mice by cardiac puncture and put in EDTA/Heparin Blood collection tubes. 100 μL of the blood was put in 1 mL of BacT/ALERT FA plus aerobic media and incubated for 1-7 days in an incubator maintained at 37 °C with 5% CO_2_. 100 μL of the samples were taken out from bacterial culture media aseptically and plated on blood agar to detect the growth of bacteria. The limit of detection of the blood culture media was determined by culturing a known CFU dilution of *S. xylosus* and *Enterobacter cloacae* strain in similar conditions. We found that the culture can detect as low as 5 CFU bacteria.

### Vagus nerve and spinal cord culture

Mice were euthanized by CO_2_ asphyxiation. The left and right cervical vagus nerves running along the neck were dissected under a dissection microscope and placed in sterile screw cap tubes with 1.5 mm ceramic beads and 500 μL of PBS. Similarly, the spinal cord was isolated from the trunk aseptically and put in the screw cap tubes. The samples were homogenized in a bead homogenizer at a speed of 5 m/s for 20 seconds twice. 200 μL of the homogenate was plated on a blood agar and incubated in an incubator maintained at 37°C with 5% CO_2_. Bacteria colonies isolated from tissue homogenates were identified by VITEK MS (Clinical Microbiology Laboratory, Emory University Hospital).

### Brain and other organ culture

Mice were euthanized by CO_2_ asphyxiation. The brain, heart, lungs, kidneys, liver, spleen, and ileum tissues were collected under aseptic conditions. Different sets of tools were kept for different organs. After each use, the instruments were sterilized by dipping in 70% ethanol and washing them in sterile PBS. The organs were collected in plastic-capped sterile glass culture tubes with 1 ml of sterile 1X PBS. The glass tubes were weighed before and after the organ collection to calculate the weights of the organs. The organs were homogenized, and the probe of the homogenizer was washed 4 times (2 washes with sterile water, 1 wash with 70% ethanol, and 1 wash with sterile PBS) in between the samples. 150-500 μL of the homogenate was plated on a blood agar and incubated in an incubator maintained at 37°C with 5% CO_2_.

### Cervical vagotomy

Mice were anesthetized using 4% isoflurane, and the surgical plane was maintained with 1.5% isoflurane. 1mg/kg of buprenorphine E.R. was injected subcutaneously as a presurgical analgesics. Mice were placed in the supine position, and the neck area was shaved and cleaned with alcohol and betadine solution. 2 cm incision was made in the skin middle of the neck using a scalpel. Under a dissecting microscope, the connective tissue layer was separated using a pair of forceps to expose mandibular glands. The connective tissue layer was further separated to expose the sternohyoid, sternomastoid, and omohyoid muscles. The sternomastoid muscle of the right side was retracted to visualize the pulsating carotid artery and the right vagus nerve. The right vagus nerve was carefully separated from the carotid sheath and a portion of the vagus nerve was cut with the help of micro-dissecting scissors. The skin was sutured, and mice were allowed to recover. SHAM surgery was performed, where the right vagus nerve was separated from the carotid sheath but not cut. Of note, bilateral cervical vagotomy was not possible since this would be fatal.

### Congo red staining

Congo red staining was performed as described previously <u>^21^</u>. Briefly, Formalin-fixed paraffin-embedded brain coronal sections were dewaxed using xylene and decreasing ethanol gradients. The sections were equilibrated in alkaline saturated NaCl solution in 80% ethanol for 20 minutes, followed by incubation in alkaline 0.2% Congo red solution for 30 minutes. The Congo red-stained slides were washed once with 95% ethanol and twice with 100% ethanol, followed by incubation in xylene. The coverslips were mounted on the slide using DPX mountant. Bright-field images were acquired in a microscope. The number and percentage area covered by the plaques were quantified using ImageJ software.

### Immunofluorescence staining

Immunofluorescence staining was performed as described previously <u>^22^</u>. Briefly, paraffin-embedded formalin-fixed brain coronal sections were dewaxed thrice in xylene, followed by xylene: ethanol (1:1). The sections were rehydrated in decreasing ethanol gradients (from 100% to 70%) and then in distilled water. The heat antigen retrieval was performed in 1X Diva decloaker buffer in a vegetable steamer. After the slides were cooled down to room temperature, the slides were washed thrice with PBS and then the permeabilization was performed in PBS/gelatin/Triton X buffer 0.25% for an hour at room temperature. The slides were blocked with 5% BSA in the permeabilization buffer. The primary antibody was diluted in 1% BSA and incubated overnight at 4•C. Slides was washed thoroughly in PBS, followed by e secondary antibody incubation for 1 hour at room temperature in 1% BSA. The coverslips were mounted using a mounting media containing DAPI and sealed with nail polish. The immunofluorescence images were acquired in Nikon A1RHD confocal microscope. ImageJ was used for image processing and quantification.

### Microglial ramification index

The ramification index of microglia was determined from the immunofluorescence images from IBA-1-stained brain coronal sections. Maximum intensity projection images of 10 μm thick sections were taken to identify microglia. The microglia ramification index is defined as the ratio of cell area divided by the projection area <u>^23-25^</u>, considering that resting microglia have thin and long projections while the inflamed and activated microglia have less ramified and retracted processes. The cell area and projection area were calculated using ImageJ software by drawing the boundaries manually around the microglia. The ramification index of a minimum of 20 microglia from a minimum of 3 mice was calculated.

### Enterobacter cloacae gavage and detection

*Enterobacter cloacae* isolate RS was described previously <u>^26^</u>. Overnight culture of *E. cloacae* (0.5 x10^7^ cfu) was orally gavaged into C57BL/6 mice previously treated with a cocktail of antibiotics. Following gavage, mice were subsequently fed Paigen diet. The colonization of *E. cloacae* was detected in the gut on day 5 after gavage. Bacteria colonies isolated from feces and ileum tissue homogenates were identified by VITEK MS as described above.

### 16S genomic sequencing

Fecal samples were collected from *Mdr2^-/-^* mice on the HFD or control diet or HFD plus antibiotics, and mouse strains BTBR (i.e., ASD), LRRK2 (i.e., PD) and APP/PS1 (i.e., AD) for DNA isolation using Qiagen Powersoil Pro DNA extraction method. Libraries were prepared using the 16S metagenomic sequencing library preparation. In this workflow, 16S amplicons were generated using KAPA HiFi HotStart ReadyMix and primers specific to the 16S (V3-V4) region. After PCR clean-up, 16S libraries were pooled in equal amounts based on fluorescence quantification. Final library pools were quantitated by quantitative PCR and sequenced on an Illumina miSeq using miSeq v3 600 cycle chemistry. Sequence analyses were performed using the NovaSeq platform. Raw data files in FASTQ format were subjected to reads merge by overlapping sequences, data quality control, and chimera filtering, resulting in high-quality clean data. Divisive Amplicon Denoising Algorithm was used for construction of Operational Taxonomic Units for analyses including diversity, taxonomy, and differential analyses.

## Acknowledgements

We thank Dr. Ralph Norgren for insightful discussions, advice, and critical review of the manuscript. We thank the Flow Cytometry, Virology and Pathology Cores of Emory Vaccine Center, and the veterinary and animal care staff of Emory National Primate Research Center, Emory University for their assistance. This study was supported in part by the Emory Integrated Genomics Core (EIGC), which is subsidized by the Emory University School of Medicine and is one of the Emory Integrated Core Facilities. We are also grateful to Lyra M. Griffiths (EIGC) for her assistance in metagenomic sequencing and analyses, and Evan Dessasau (Pathology Core) for their assistance in histopathological analyses. We are also thankful to LC Sciences (Houston) Bioinformatics services for their assistance in analyses including diversity, taxonomy, and differential analyses for bacterial families.

## Data availability statement

All data are available in the main text or the supplementary materials. All data, code, and materials used in the analysis will be available to any researcher for purposes of reproducing or extending the analysis.

