## Supplemental Figures 1-4 for "Translocation of gut commensal bacteria to the brain"

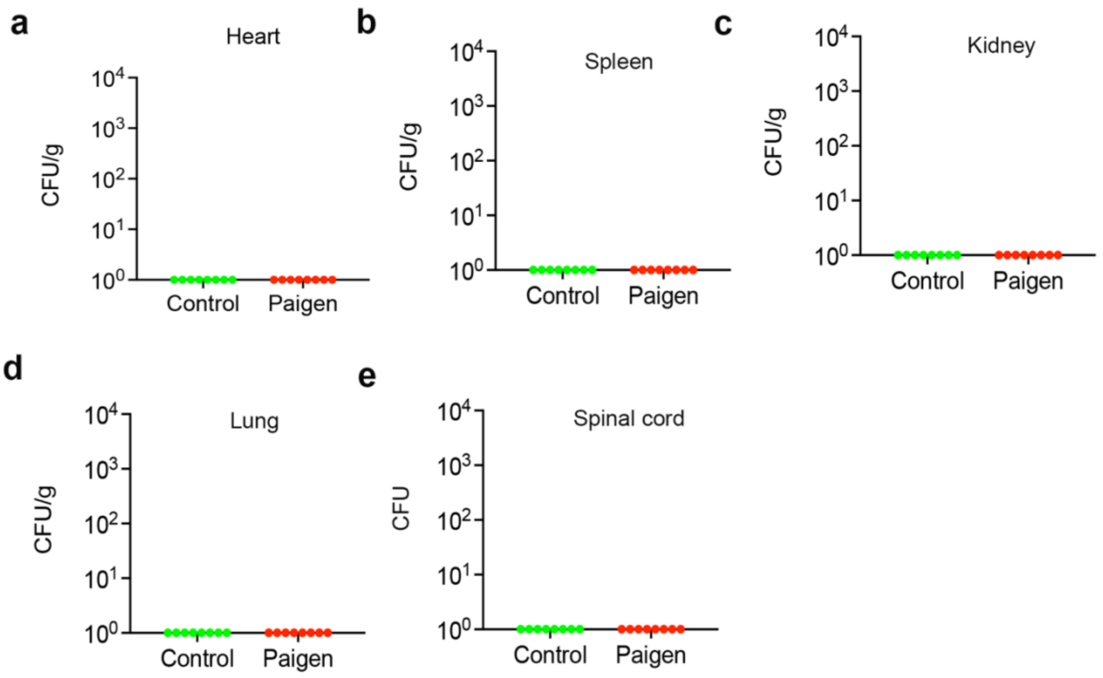


**Supplementary Figure S1**: Total colony forming units (CFU) of bacteria (a) per gram of heart, (b) per gram of spleen, (c) per gram of kidney, (d) per gram of lungs, and (e) the spinal cord are shown from Mdr2-/- mice fed with either regular diet (control) paigen diet (paigen).

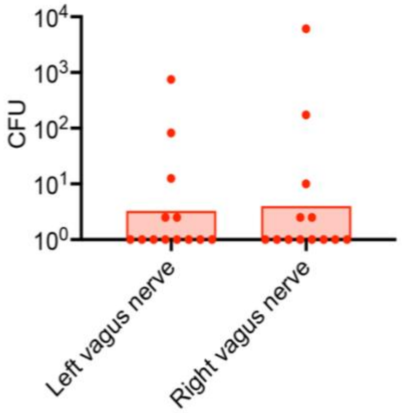


**Supplementary Figure S2**: Total colony forming units (CFU) of bacteria in the left and right vagus nerve from Mdr2-/- mice fed with paigen diet.

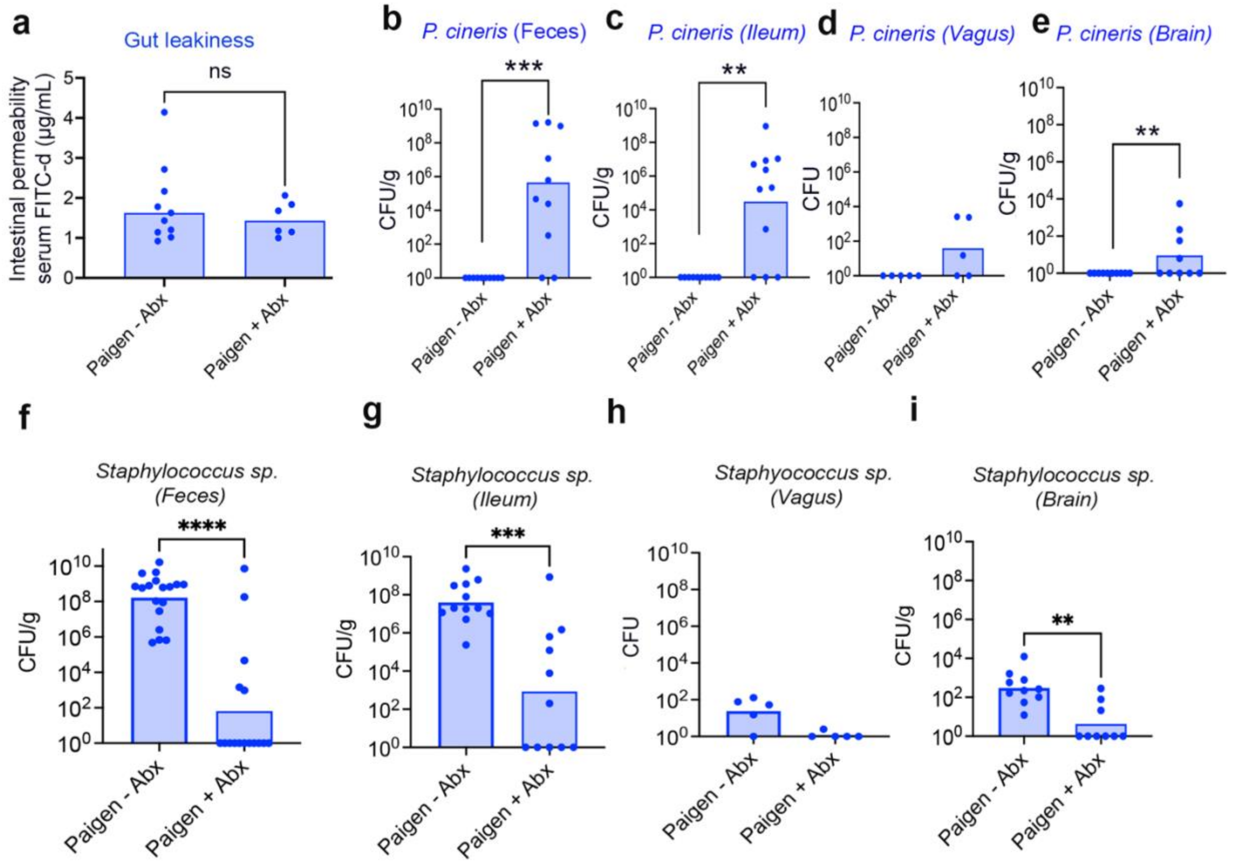


**Supplementary Figure S3**: 12-week-old Mdr2-/- mice were treated with a cocktail of antibiotics during Paigen diet feeding. a) Intestinal permeability in antibiotics-treated (Abx) Mdr2-/- mice were determined. b-e) CFU of *Paenibacillus cineris* in (b) feces, (c) ileum, (d) vagus nerve, and (e) brain of antibiotic-treated (Abx) mice are shown. f-i) CFU of *Staphylococcus sp.* in (f) feces, (g) ileum, (h) vagus nerve, and (i) brain of antibiotic-treated (Abx) mice are shown.

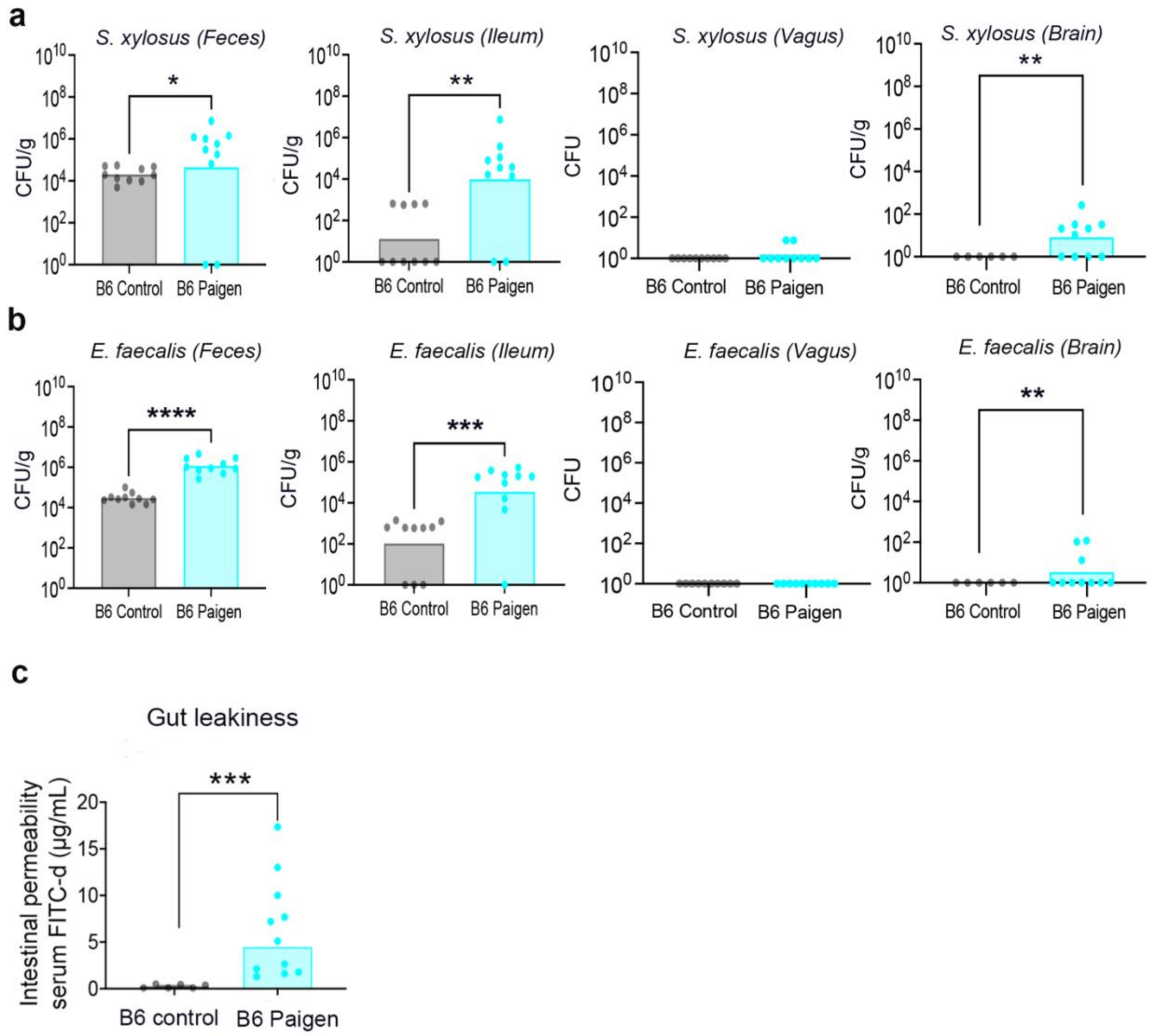


**Supplementary Figure S4**: a) CFU of *S. xylosus* in feces, ileum, vagus nerve and brain of C57BL/6 (B6) mice fed Paigen diet are shown. b) CFU of *E. faecalis* in feces, ileum, vagus nerve and brain of C57BL/6 (B6) mice fed Paigen diet are shown. c) Intestinal permeability in Paigen diet-fed B6 were determined.
